# Spatial Transcriptomics of Schizophrenia Insular Cortex Reveals Blood-Brain Barrier Hyperglycolysis and Increased Parenchymal Mitochondrial Respiration

**DOI:** 10.1101/2025.03.12.642911

**Authors:** K. Huizer, V. van Dis, T. Tran, P.R. Bakker, M. Claesen, I. Nijman, R. Geene, N.J.M van Beveren, D.A.M. Mustafa

**Author notes:** These authors contributed equally to this work (K. Huizer, V. van Dis, T. Tran, D.A.M. Mustafa). Correspondence to: dr. K. Huizer / dr. D.A.M. Mustafa /.

## Abstract

**Introduction:** The blood-brain barrier (BBB) acts as the metabolic and immunological gatekeeper of the brain. Since alterations in neurometabolism and neuroimmunity are found in schizophrenia-spectrum disorders (SSD) which are hypothesised to be important disease mechanisms, we aimed to investigate whether changes in BBB function could underly these findings using a novel spatially resolved transcriptomics technique.

**Methods:** Formalin-fixed paraffin-embedded insular cortex tissue from 8 brain donors with SSD and 8 matched controls derived from the Netherlands Brain Bank-Psychiatry were selected for whole transcriptome analysis (GeoMx Human Whole Transcriptome Atlas) on the GeoMx Digital Spatial Profiler platform. Combining nuclear staining with an endothelial cell marker (CD31) allowed for the separation of BBB and parenchyma areas of interest (AOIs) for downstream sequencing on the Illumina NextSeq 2000. For each sample, biological triplicates were sequenced. Comparing SSD to control for both the BBB and parenchyma AOIs, differentially expressed genes (DEGs) were identified using a Linear Mixed Model, a heatmap was created displaying all genes with a false-discovery rate <0.01, and Fast Gene Set Enrichment Analysis was used for pathway analysis.

**Results:** A total of 96 whole transcriptome profiles were generated (24 BBB and 24 parenchyma for both SSD and controls). Expression of endothelial genes (*PECAM1/CD31, CLDN5, VWF, CD34, ENG*) was significantly increased in BBB, confirming enrichment of endothelial cells (ECs) in this AOI. Cluster analysis showed perfect clustering of BBB versus parenchyma, and good clustering of SSD samples within the BBB cluster. At a |Log2FC| ≥ 0.25, we identify 265 significantly DEGs in the BBB AOI and 6 in the parenchyma AOI comparing SSD to control. Pathway analysis revealed a distinct metabolic transcriptional profile in SSD, characterized by hyperglycolysis in the BBB and increased mitochondrial energy metabolism in parenchyma.

**Conclusion:** Our findings implicate the BBB in the metabolic pathophysiology of SSD. Furthermore, our findings add nuance to the existing understanding of brain bioenergetic alterations in SSD, suggesting that metabolic changes may be region-specific rather than generalized. This highlights the need for a ‘brain mapping’ approach examining multiple brain regions from the same donor. Finally, the distinct metabolic profiles of the BBB and brain parenchyma emphasize the importance of spatial multi-omics in post-mortem psychiatric research and the potential for therapies targeting BBB function in SSD.

## Introduction

Within the central nervous system, neurons need a suitable microenvironment to safeguard synaptic communication. The blood-brain barrier (BBB) — a highly evolved structure consisting of brain endothelial cells (ECs), pericytes, a basal membrane, and astrocyte end-feet^1^ — separates the blood from the brain tissue proper (the parenchyma). The BBB tightly and dynamically regulates which cells and substances enter and exit the brain, and secretes signalling factors to safeguard homeostasis. The BBB thus functions as a dynamic physical, transport, metabolic and immunological barrier. As such, the BBB is a crucial regulator of the neuronal microenvironment. The concept of the neurovascular unit describes how neurons and astrocytes signal their homeostatic needs to the adjacent BBB, e.g., by increasing the supply of energy substate when neuronal activity is high^2,3^. Therefore, the BBB is a functionally heterogeneous structure depending on its location in the vascular tree (e.g., arteriole, venule, capillary), its anatomic brain location (e.g., thalamus versus prefrontal cortex), but also based on the dynamic needs of the tissue it supplies (e.g., high vs low neuronal activity)^2,3^. Since the largest surface area of blood vessels in the brain consists of capillaries, the BBB in the brain capillary bed is quantitatively the most important for brain homeostasis^4^.

In the last decade, converging evidence points towards immune and metabolic aberrations as contributing factors to schizophrenia-spectrum disorders (SSD)^5,6^. As the metabolic and inflammatory gatekeeper to the brain, malfunction of the BBB could explain these findings. Importantly, involvement of the BBB in schizophrenia could have clinical implications since the BBB is therapeutically easily accessible through the blood. The hypothesis that the BBB could be implicated in schizophrenia is only recently being investigated^7,8^. Its small size and close integration with the brain parenchyma has made it very challenging to reliably study the BBB. Imaging techniques (e.g., fMRI and MRS) do not offer sufficient resolution to study the BBB at the (sub)cellular level. Postmortem brain research can provide the molecular resolution required^9,10^, but thus far depended on tissue homogenisation to allow for whole transcriptome analysis, rendering a separation between the BBB and brain parenchyma impossible. Recently, novel techniques have been developed allowing for whole transcriptome analysis while retaining crucial spatial information^11,12^, therefore theoretically allowing for the omics-scale analysis of the BBB. However, at present, spatially resolved transcriptomics has predominantly been used in cancer research^13,14^. Characteristically, tumour tissue exhibits large transcriptional changes compared to healthy tissue and can be easily morphologically separated. In contrast, the molecular biology of psychiatric disorders is heterogeneous, characterized by small transcriptional changes in many genes in morphologically normal appearing tissue^7,15^. Another caveat to spatial transcriptomics of the BBB is the limited number of BBB cells within the maximum field of analysis compared to for instance tumour tissue. Additionally, for any post-mortem human brain study, the usual confounding factors including post-mortem delay, fixation time, agonal changes, cause of death, comorbidities etc. need to be taken into account^16–20^ To further complicate this research field, psychiatric diagnoses are syndromal, with overlapping diagnostic criteria; diagnoses as such do not (yet) reflect clear-cut differences in underlying pathophysiology^21^. Furthermore, there is a general shortage of donated brains from psychiatric patients^18,22^, especially of schizophrenia-spectrum disorder (SSD) patients, often leading to added issues regarding tissue quality and sample quantity. Altogether, the technical and practical feasibility of using spatial transcriptomics to study SSD in general, and the BBB in particular, remains to be determined.

The current study uses spatially resolved whole transcriptome analysis on the BBB and on the surrounding brain parenchyma in post-mortem insular cortex tissue derived from donors with SSD compared to non-affected controls, using Nanostring’s GeoMx Digital Spatial Profiling, Human Whole Transcriptome Atlas. We selected insular cortex tissue in our study because, despite its clear involvement in SSD^23–28^, this area remains understudied^23,24^.

## Materials and methods

### Institutional Review Board Approval

The study was approved by the Erasmus MC Institutional Review Board (METC) under the number *MEC-2023-0373*. The Review Board confirmed that the rules laid down in the Medical Research Involving Human Subjects Act (also known by its Dutch abbreviation WMO), do not apply to this research proposal.

### Tissue Samples

Formalin-fixed paraffin-embedded (FFPE) post-mortem brain samples of the insular cortex were selected for this study. The clinical inclusion criteria included a main diagnosis of schizophrenia or schizoaffective disorder for the SSD cohort and the absence of a psychiatric diagnosis for the control cohort, with an age range of 18–80 years. Notably, SSD samples were matched to controls as closely as possible for gender, age, and overall cause of death (e.g., cancer, cardiovascular, respiratory, euthanasia) to create a homogeneous sample set. Exclusion criteria included intracerebral pathology such as malignancy, cerebrovascular accidents affecting the insular cortex, neuro-inflammatory diseases, and neurodegeneration (Braak stage > 2). Additionally, to minimize the confounding effects of post-mortem delay (PMD) and low pH on gene expression analyses^18,29^, we strived to include samples with a PMD < 24 hours and a post-mortem cerebrospinal fluid (CSF) pH ≥ 6.

For spatial transcriptomics analysis, additional technical inclusion criteria were applied: FFPE samples with a maximum fixation time of 8 weeks, samples stored as FFPE tissue blocks for a maximum of 15 years, and freshly cut 5 um tissue sections prepared and stored at 4°C for a maximum of 3 days before analysis.

The Netherlands Brain Bank – Psychiatry (NBB-Psy; https://www.brainbank.nl) exclusively stores brain donor material with a PMD < 20 hours, typically <12 hours. NBB-Psy follows a standardized protocol for donor brain preparation, including a formalin fixation time of 4 weeks and the storage of extensively sampled brain tissue in both FFPE and fresh-frozen formats. Additionally, NBB-Psy provides comprehensive, high-quality clinical information on donor medical and psychiatric history, cause of death, agonal state, and a standardized neuropathological examination. Based on these well-defined standards, we proceeded with material from NBB-Psy.

### Digital Spatial Whole Transcriptome Atlas Profiling

We used the GeoMx Human Whole Transcriptome Atlas^30^ (WTA) on the GeoMx Digital Spatial Profiler (DSP) Platform^31^ to measure our samples following the manufacturer’s instructions^11,32^ (Nanostring®, a Brucker company; Seattle). Slide preparation was performed on the Leica Bond RXm system according to the manufacturer’s instructions and as previously described by Merritt et al.^11^. Freshly cut 5 μm FFPE tissue sections were loaded onto the Leica Bond RXm system, where they underwent baking, deparaffinization, and epitope retrieval using ER2 solution (Leica Biosystems, AR9640) for 20 minutes at 100°C. This was followed by incubation for 15 minutes at 37°C with 1 μg/mL proteinase K (Thermo Fisher Scientific, AM2546) and post-fixation in 10% neutral buffered formalin (NBF). The slides were then incubated overnight at 37°C with the Human Whole Transcriptome Atlas (WTA) probe panel. After washing in 50% formamide at 37°C, blocking was performed, followed by immunostaining with CD31 (Abcam, ab215912) and with SYTO13 nuclear staining (Thermo Fisher Scientific, S7575). After staining, the slides were loaded and scanned on the GeoMx instrument.

### Defining Regions of Interest (ROIs) and Areas of Interest (AOIs)

Morphological markers (CD31 and nuclear staining) were used to select Regions of Interest (ROIs) in insular cortical layers III-VI containing most microvessels. Selection was performed under the guidance of an experienced neuropathologist (VvD), and with help by an experienced technician (RG) as follows: 1) CD31+ (BBB): microvessels positive for both CD31 and nuclear staining; 2) CD31-(parenchyma): brain parenchyma adjacent to the BBB, negative for CD31 but positive for nuclear staining (Fig. 1). For the BBB AOI, settings were adjusted to include cells bordering on CD31+ cells, to enrich for pericytes and astrocyte end-feet in direct contact with microvascular endothelial cells, thus capturing the entire BBB. Biological triplicates were selected for each AOI within each tissue section, resulting in 6 unique whole transcriptome profiles per sample (3 BBB AOIs and 3 parenchyma AOIs per sample). In total, this yielded 96 WTA profiles. The recommended minimum of 100 nuclei/AOI could not always be achieved in the BBB AOI. Therefore, the minimum nuclear count per AOI was adjusted to 40 nuclei, which was met for all AOIs. For the BBB (CD31+) AOIs, the average nuclear count was 86 (lowest 47, highest 171). For the Parenchyma (CD31-) AOIs, the average nuclear count was 1142 (lowest 220, highest 5256). See **Figure 1**.

**Figure 1:**
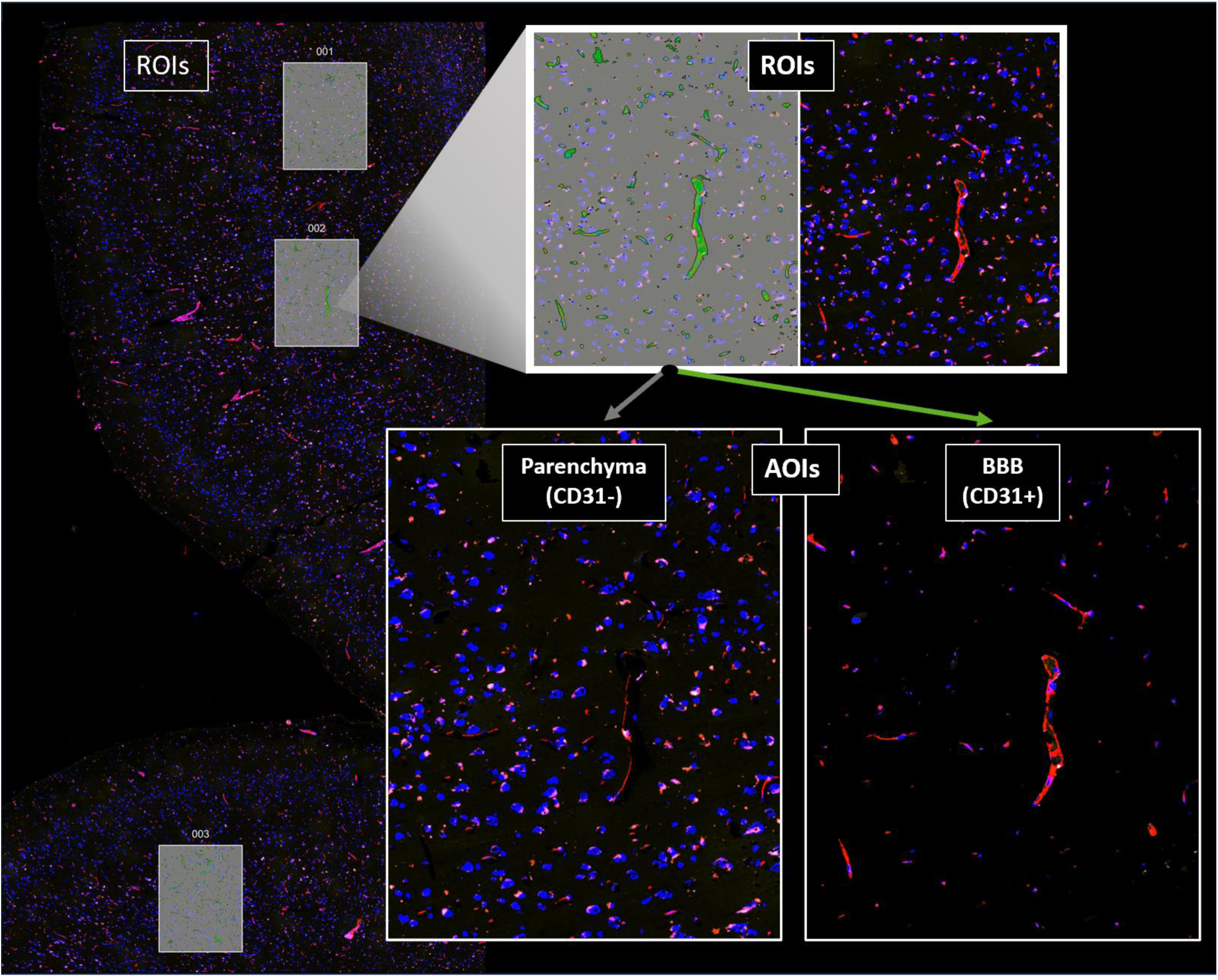
Example of Region of Interest (ROI) and Area of Interest (AOI) selection. The image provides a representative example of the selection process of ROIs (left of image, small magnification: rectangles 1, 2 and 3) and AOIs (blow-up of rectangle 2; right bottom figure, high magnification) in insular cortex samples. For each ROI, 2 AOIs were selected: CD31+ Blood-brain barrier (BBB) and CD31-parenchyma. In total, 6 AOIs per tissue sample were analysed with whole transcriptome analysis (3x BBB, 3x parenchyma).Visual inspection of the slides (VvD) revealed that most background staining was due to neuronal lipofuscin. Therefore, within the BBB (CD31+) AOI, cells with an aspecific fluorescent signal in other channels were excluding from the AOI, ensuring that background signal was removed. Within the parenchyma (CD31-) AOI, aspecific fluorescent signal was included to avoid exclusion of lipofuscin-containing neurons.

### Sequencing

Sequencing libraries were prepared using the GeoMx seq code Kit. Amplified libraries were pooled and purified using AMPure XP beads (Beckman Coulter, A63880). Sequencing was performed on an Illumina NextSeq 2000, using a P3 50-cycle kit with a paired-end 2×27 bp configuration. Raw sequencing data were demultiplexed and data processing was conducted using the GeoMx NGS Pipeline.

### Data Analysis

The WTA profiles were compared between SSD and controls for both BBB and parenchyma AOIs. Analysis was performed based on a standardized workflow as recommended by NanoString/Bruker (https://bioconductor.org/packages/release/bioc/html/GeomxTools.html) but extended by USEQ (<u>UMCUGenetics/GeoMx-DSP-analyses: Useq GeoMX DSP pipeline revision</u>).

Differentially expressed genes were identified using a linear-mixed model (LMM), corrected for multiple testing^33^. A heatmap was created for all genes with an FDR < 0.001 in the LMM analysis. LMM results also served as input for pathway analysis using Fast Gene Set Enrichment Analysis (FGSEA).

## Results

### Description of sample characteristics

Based on our clinical and technical criteria described above, we selected insular cortex samples from 8 donors with SSD and 8 control donors. See **Table 1** for demographic, clinical and technical sample characteristics.

**Table 1:** Demographic, clinical and technical sample characteristics.

| nbb | Psychiatric Diagnosis | gender | age | Cause of Death | Post-Mortem Delay (h:m) | Braak stage | Ph liquor |
| --- | --- | --- | --- | --- | --- | --- | --- |
| 2010-021 | schizophrenia | M | 59 | Cardiac arrest with renal insufficiency and infection | 12:30 | ? | 5.93 |
| 2010-055 | schizophrenia | M | 64 | Pulmonary embolism | 19:15 | 1 | 7 |
| 2010-127 | schizoaffective disorder, depressive type | F | 79 | Heart fibrillation with heart failure | 4:45 | 1 | 6.34 |
| 2012-031 | schizoaffective disorder, depressive type | F | 55 | Euthanasia | 9:50 | 0 | 6.82 |
| 2016-003 | schizophrenia | M | 67 | Euthanasia for terminal gastro-esofagal cancer | 5:45 | 1 | 6.29 |
| 2018-102 | schizophrenia | F | 65 | Cachexia and dyspnea due to metastasised terminal breast cancer | 7:10 | 2 | 6.4 |
| 2021-62 | schizoaffective disorder, bipolar type | M | 59 | Metastatic kidney cancer (bone, pulmonary, lymphatic) | 4:25 | 1 | 6.71 |
| 2022-115 | schizophrenia | F | 80 | Old age, renal insufficiency | 8:20 | 2 | 6.79 |
| 2011-072 | none | F | 76 | Hepatic failure and colon carcinoma | 7:15 | 2 | 6.87 |
| 2012-049 | none | F | 70 | Cachexia due to endstage pancreas carcinoma | 7:35 | 2 | 6.03 |
| 2012-059 | none | F | 78 | Bronchopneumonia with neuroendocrine carcinoma pancreas, metastasised | 4:35 | 2 | 6.41 |
| 2012-104 | none | M | 79 | Euthanasia (terminal heartfailure with dyspnoea) | 6:30 | 2 | 6.71 |
| 2014-043 | none | F | 60 | Metastasized mammacarcinoma | 8:10 | 0 | 6.58 |
| 2018-121 | none | F | 68 | Acute infection with metastatic pancreatic cancer | 3:30 | 0 | 7.37 |
| 2021-051 | none | M | 77 | Metastatic prostate cancer | 7:25 | 1 | 6.72 |
| 2021-119 | none | M | 73 | Collapse, followed by asystole. Probably cardiac death | 10:15 | 1 | 6.71 |

### Areas of Interest (AOIs) And Gene Detection

A total of 96 AOIs were selected and sequenced across all samples, including 24 BBB (CD31+) and 24 parenchyma (CD31-) AOIs in both the SSD and the control group. Transcriptomic sequencing data from all AOIs, except one, passed the QC step. In total, 7,321 genes passed the identification threshold and were remained for analysis.

### Successful Selection of BBB AOIs

To validate the accuracy of the BBB (CD31+) and parenchyma (CD31-) AOI selection, we assessed if endothelial cell specific gene expression was enriched in BBB. Five genes known to be highly and specifically expressed in endothelial cells were selected (*PECAM1/CD31, CLDN5, VWF, CD34, ENG*). Their expression levels were compared between the BBB and parenchyma AOIs. All endothelial cell genes showed a significantly higher expression in the BBB compared to parenchyma, confirming enrichment of endothelial cells within the BBB AOI (**Figure 2**).

**Figure. 2:**
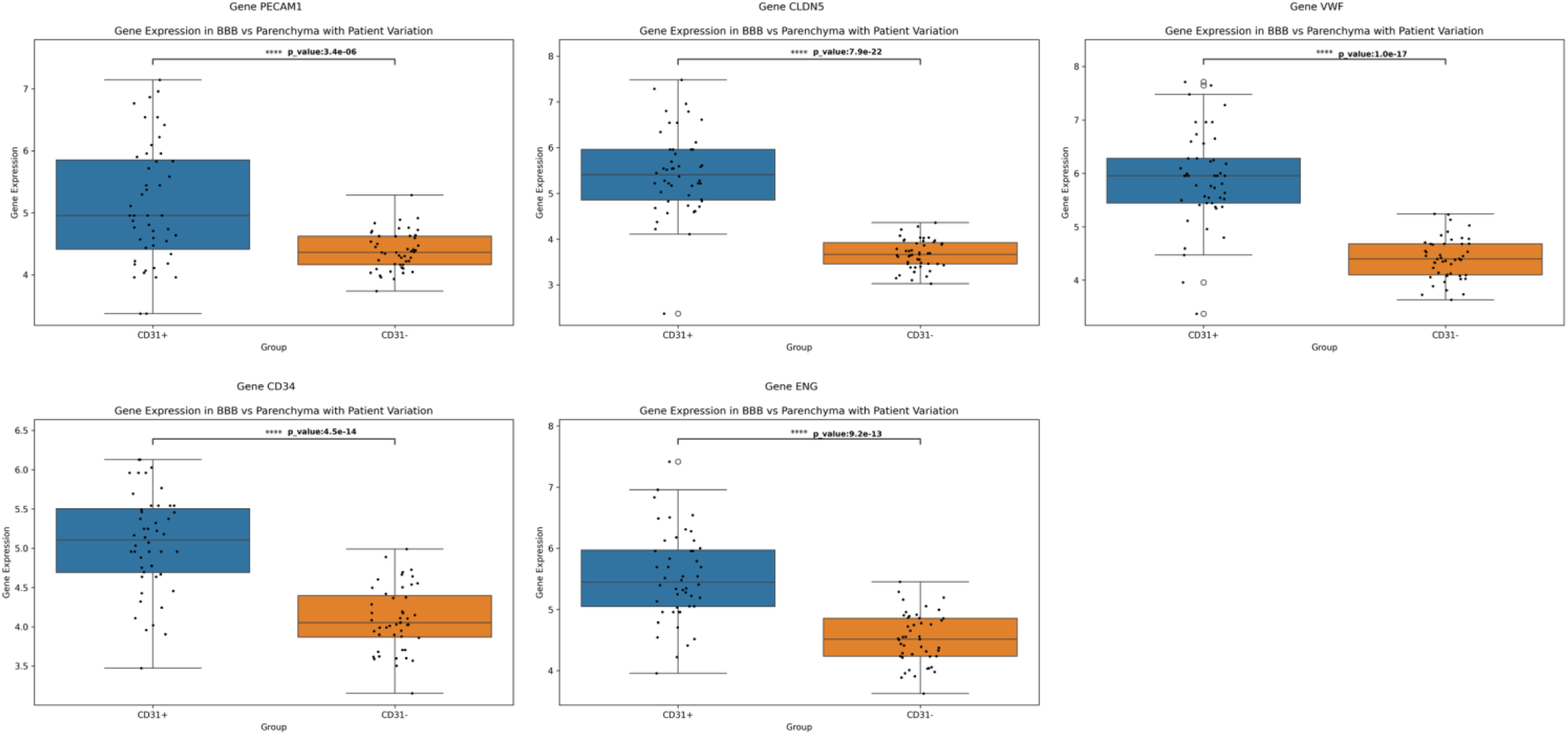
Boxplots of gene expression levels of endothelial cell markers in BBB and parenchyma AOIs. To examine whether our technical setup enriched for endothelial cells in the BBB AOI, we generated boxplots showing the gene expression levels of various endothelial cell markers (*PECAM1/CD31, CLDN5, VWF, CD34, ENG*). As can be seen in the figure, all markers were significantly enriched in the BBB AOI compared to the parenchyma AOI.

### Differentially Expressed Genes in BBB (CD31+) between SSD and Control

Comparing the gene expression profiles of BBB AOIs in SSD versus controls using LMM, we identified six differentially expressed genes (DEGs) at an absolute Log2 Fold Change (|Log2FC|) cutoff of ≥1 and corrected p ≤0.05. All 6 genes were downregulated in the BBB of SSD compared to controls (**Table 2**).

**Table 2:** BBB DEGs between Control and SSD with a|Log2FC| ≥ 1.0 and a corrected p-value ≤ 0.05. A positive Log2FC indicates higher expression in control vs SSD.

| Gene | Log2FC | corr. P-value |
| --- | --- | --- |
| TNFRSF6B | 1.37 | 0.01 |
| IGFBP3 | 1.19 | 0.00 |
| DRC3 | 1.08 | 0.03 |
| ARHGAP27 | 1.06 | 0.01 |
| FAM210A | 1.03 | 0.01 |
| P4HA1 | 1.02 | 0.02 |

However, post-mortem brain whole transcriptome studies in psychiatry employ a less stringent |Log2FC| cutoff value (≥ 0.25)^7,15^, to account for the more subtle transcriptional differences in psychiatric diseases as compared to diseases like cancer. At a |Log2FC| ≥ 0.25, we identify 265 significantly DEGs in the BBB AOI, including 228 downregulated and 37 upregulated in SSD compared to control.

### Limited Differentially Expressed Genes in Parenchyma (CD31-)

In parenchyma, applying the stringent |Log2 FOC| threshold of ≥1 yielded no significantly DEGs between SSD and control. Adjusting the |Log2 FOC| cutoff to ≥ 0.25 yielded only 6 significantly DEGs (2 downregulated and 4 upregulated in SSD compared to control). Effect sizes were therefore much smaller in parenchyma than in BBB. One DEG in parenchyma was involved in energy metabolism (COX7A1, decreased in SSD).

### Clear Clustering of BBB vs Parenchyma & Strong Clustering of SSD vs Control BBB

All genes with a false discovery rate (FDR) < 0.001 (LMM) were displayed in a heatmap using unsupervised cluster analysis (**Figure 3**). The insular cortex BBB (CD31+) and parenchyma (CD31-) AOIs clustered separately. Within the BBB cluster, a good distinction between SSD and controls can be made. Within the parenchyma cluster, no clear clustering of SSD samples was visible.

**Figure 3:**
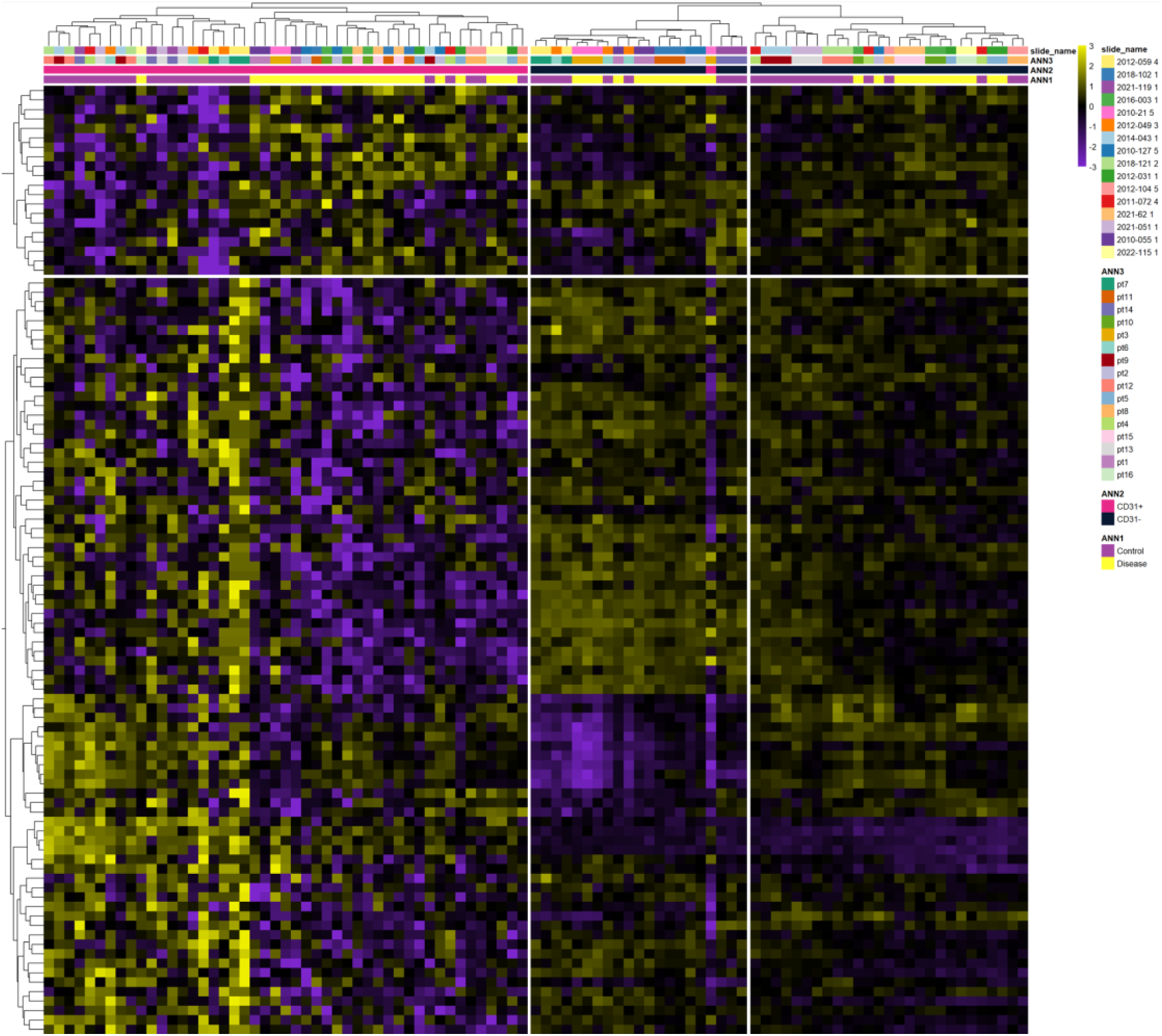
Unsupervised Cluster Analysis of all AOIs and genes with an FDR < 0.001. Heatmap of unsupervised cluster analysis of all genes with a false discovery rate (FDR) < 0.001. Yellow colour indicated increased gene expression; purple colour reduced gene expression. The first cluster separates the BBB from the parenchyma AOI. More downstream, there is a good distinction between SSD (‘disease’) and controls within the BBB cluster, but not within the parenchyma cluster.

### Distinct Metabolic Signature in SSD Insular BBB and Parenchyma: Insights from Pathway Analysis

Pathway analysis using FGSEA in both BBB and parenchyma revealed several significantly differentially enriched pathways between SSD and control. Selecting the top 15 positively and top 15 negatively enriched pathways yielded a list of in total 20 significantly differentially enriched pathways in BBB, and 20 in parenchyma (**Figure 4)**. In BBB, 6 pathways were enriched in SSD, while in parenchyma 13 pathways were enriched in SSD. Notably, pathways related to energy metabolism were disproportionately represented in both BBB and parenchyma. In SDD BBB, three glucose metabolism pathways (*Gluconeogenesis; Glycolysis; Glucose Metabolism*) were positively enriched, whereas in SSD parenchyma, seven pathways associated with mitochondrial energy metabolism (*Respiratory Electron Transport, ATP Synthesis by Chemiosmotic Coupling, and Heat Production by Uncoupling Protein; Cristae Formation; Complex I Biogenesis; TCA Cycle and Respiratory Electron Transport; Respiratory Electron Transport; Formation of ATP by Chemiosmotic Coupling; Mitochondrial Protein Import*) were positively enriched. Given this strong emphasis on energy metabolism and genes, alongside the relatively smaller effect sizes of DEG in the parenchyma AOIs, the next section will primarily explore findings related to the energy metabolism within the insular cortex BBB.

**Figure 4:**
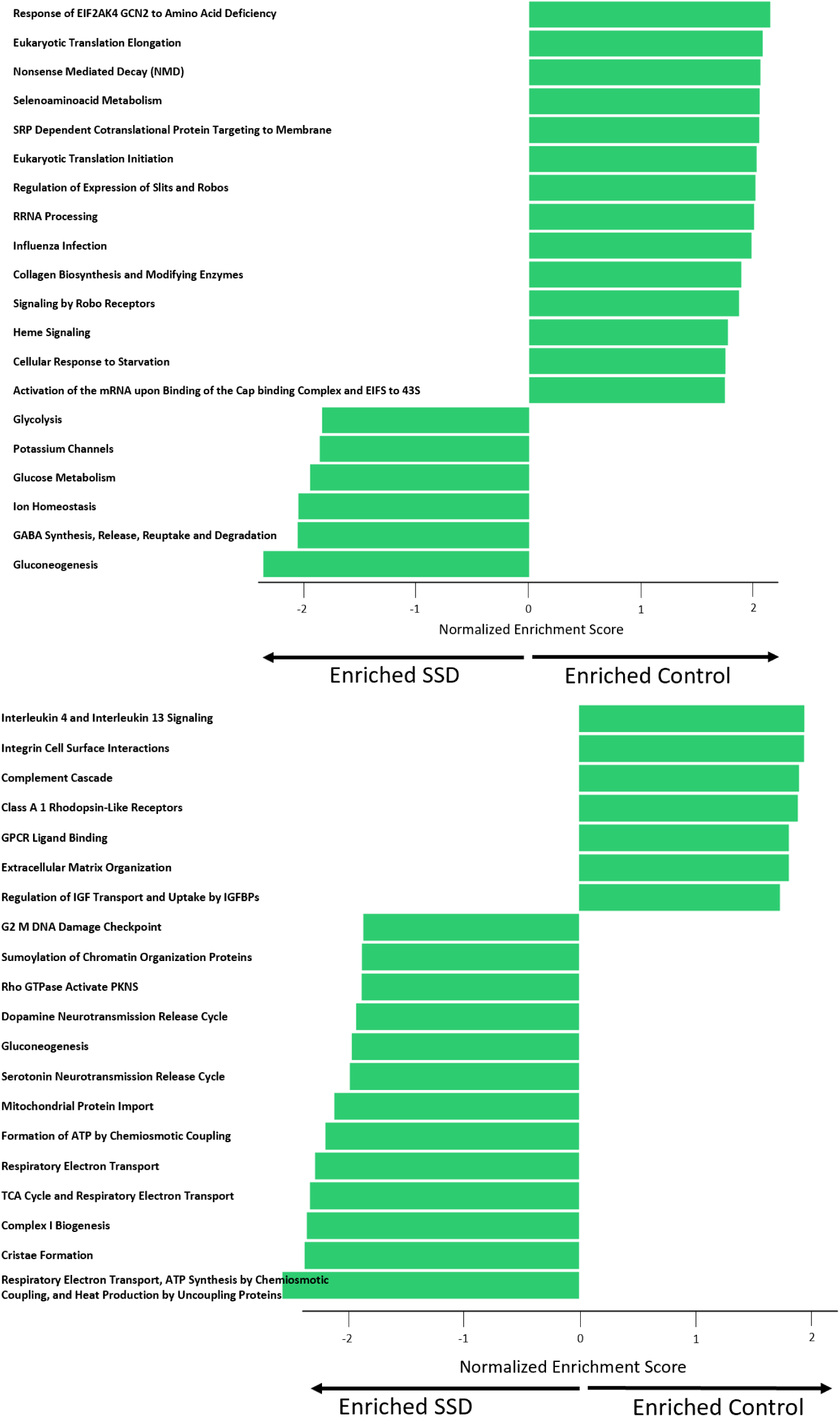
Normalised Enrichment Scores of Significantly Enriched Pathways (FGSEA) Barplots of normalized enrichment scores (NES) of significant (adjusted p-value p≤0.5) positively and negatively enriched Reactome pathways (FGSEA). A positive NES indicates enrichment of pathway genes in control vs SSD; a negative NES indicates enrichment of pathway genes in SSD vs control (<u>Top</u>: BBB; <u>Bottom</u>: Parenchyma).

Given the overrepresentation of energy metabolism pathways and genes, combined with the much smaller amount and effect sizes of DEGs in the parenchyma AOI, the next section will largely focus on our findings regarding energy metabolism the insular cortex BBB.

### Upregulation of Glycolysis in SSD BBB

To further investigate glucose metabolism in the BBB, we focused on the DEGs within the 3 enriched pathways related to glucose metabolism: *Gluconeogenesis, Glycolysis* and *Glucose Metabolism*. These three pathways shared many overlapping genes, with *Glucose Metabolism* encompassing genes from both the *Gluconeogenesis* and *Glycolysis* pathways. Therefore, we narrowed our analysis to the *Gluconeogenesis* and *Glycolysis* pathways. In our BBB dataset, 37 genes were present in the *Glycolysis* pathway and 16 genes in the *Gluconeogenesis* pathways. Seven genes were common to both pathways, encoding for enzymes that can convert substrates back and forth. These 7 shared genes were overrepresented in the top upregulated DEGs of the *Gluconeogenesis* pathway, therefore disproportionately driving the Normalized Enrichment Score (NES) for that pathway. This implies the *Gluconeogenesis* pathway is enriched in the pathway list primarily based on genes shared with the *Glycolysis* pathway. Consequently, we focused exclusively on the *Glycolysis* pathway.

Since Endothelial cells (ECs) exhibit a specialized form of glycolysis, known as aerobic glycolysis, where pyruvate is converted into lactate by lactate dehydrogenase (LDH, **Figure 5**), we included the genes coding for lactate dehydrogenase (*LDHA* and *LDHB*) to the list of DEGs in the *Glycolysis* pathway. Notably, in the *Aerobic Glycolysis* pathway in BBB, 1 gene reached significance after correcting for multiple comparisons: *PFKFB3* was upregulated in SSD BBB (Log2FC = -0.47; corrected p-value = 0.002). *PFKFB3* codes for 6-phosphofructo-2-kinase/fructose-2,6 bisphosphatase 3, which converts fructose-6-phosphate (F6P) into fructose-2,6-bisphosphate (F2,6BP). F2,6BP is a strong allosteric activator of phosphofructokinase-1 (PFK1), a rate-limiting enzyme of glycolysis^34^. Additionally, PFKFB3 is the most important driver of glycolysis in ECs^34,35^. Since the expression of *PFKFB3* is regulated by various different stimuli^34–36^, it is beyond the scope of this study to investigate the possible mechanisms behind the upregulation of *PFKFB3* in SSD BBB. See **Figure 6** for a graphic representation of DEGs in the *Aerobic Glycolysis* pathway in SSD BBB.

**Figure 5:**
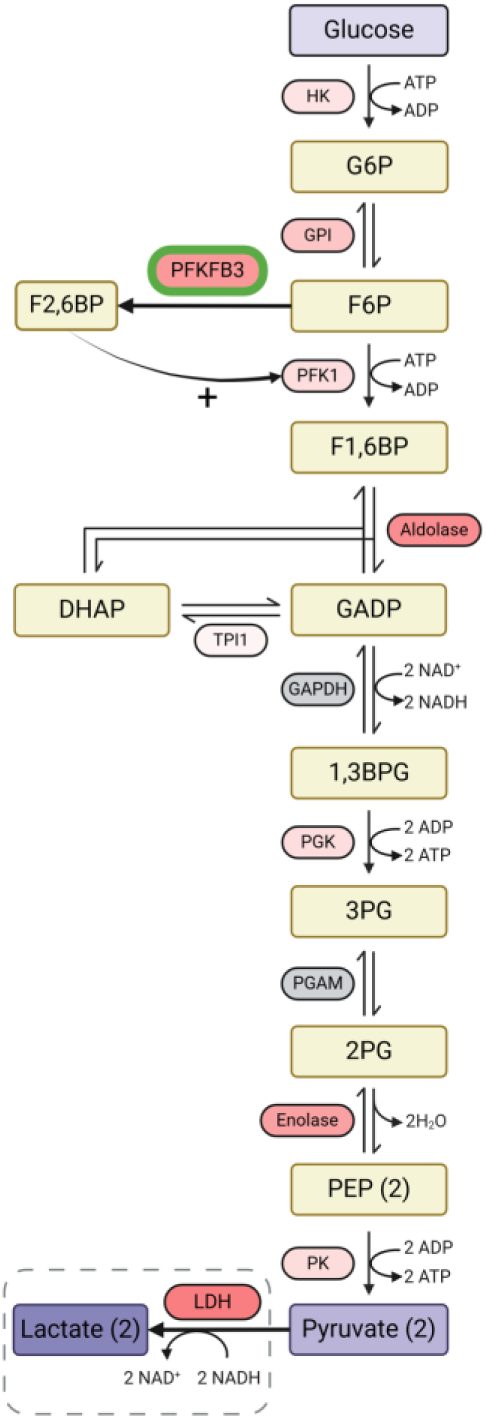
Aerobic Glycolysis Pathway in BBB. Enzymes (oval boxes) have been colour-graded based on their gene expression levels in SSD compared to control in insular cortex BBB (**red**: increased expression; deeper hues of red indicate more upregulation; **grey**: gene did not pass QC; **green stroke**: significantly upregulated at corrected p-value < 0.05). The conversion of pyruvate to lactate by LDH (dashed grey box) is not usually considered part of the *Glycolysis* pathway, but important in ECs which are almost exclusively using aerobic glycolysis. **<u>Abbreviations</u>**: 1,3BPG: 1,3-bisphosphoglycerate; 2PG: 2-phosphoglycerate; 3PG: 3-phosphoglycerate; ADP: adenosine diphosphate; ATP: adenose triphosphate; DHAP: dihydroxyacetone phosphate; F1,6BP: fructose-1,6-bisphosphate; F2,6BP: fructose-2,6-bisphosphate; F6P: fructose-6-phosphate; G6P: glucose 6-phosphate; GADP: glyceraldehyde-3-phosphate; GADPH: glyceraldehyde-3-phosphate dehydrogenase; GPI: glucose-6-phosphate isomerase; HK: hexokinase; LDH: lactate dehydrogenase; NAD: nicotinamide adenine dinucleotide; NADH: nicotinamide adenine dinucleotide; PEP: phosphoenolpyruvate; PFK1: phosphofructokinase-1; PFKFB3: 6-phosphofructo-2-kinase/fructose-2,6 bisphosphatase 3; PGAM: phosphoglycerate Mutase 2; PGK: phosphoglycerate kinase 1; PK: pyruvate kinase; TPI1: triosephosphate isomerase 1

**Figure 6:**
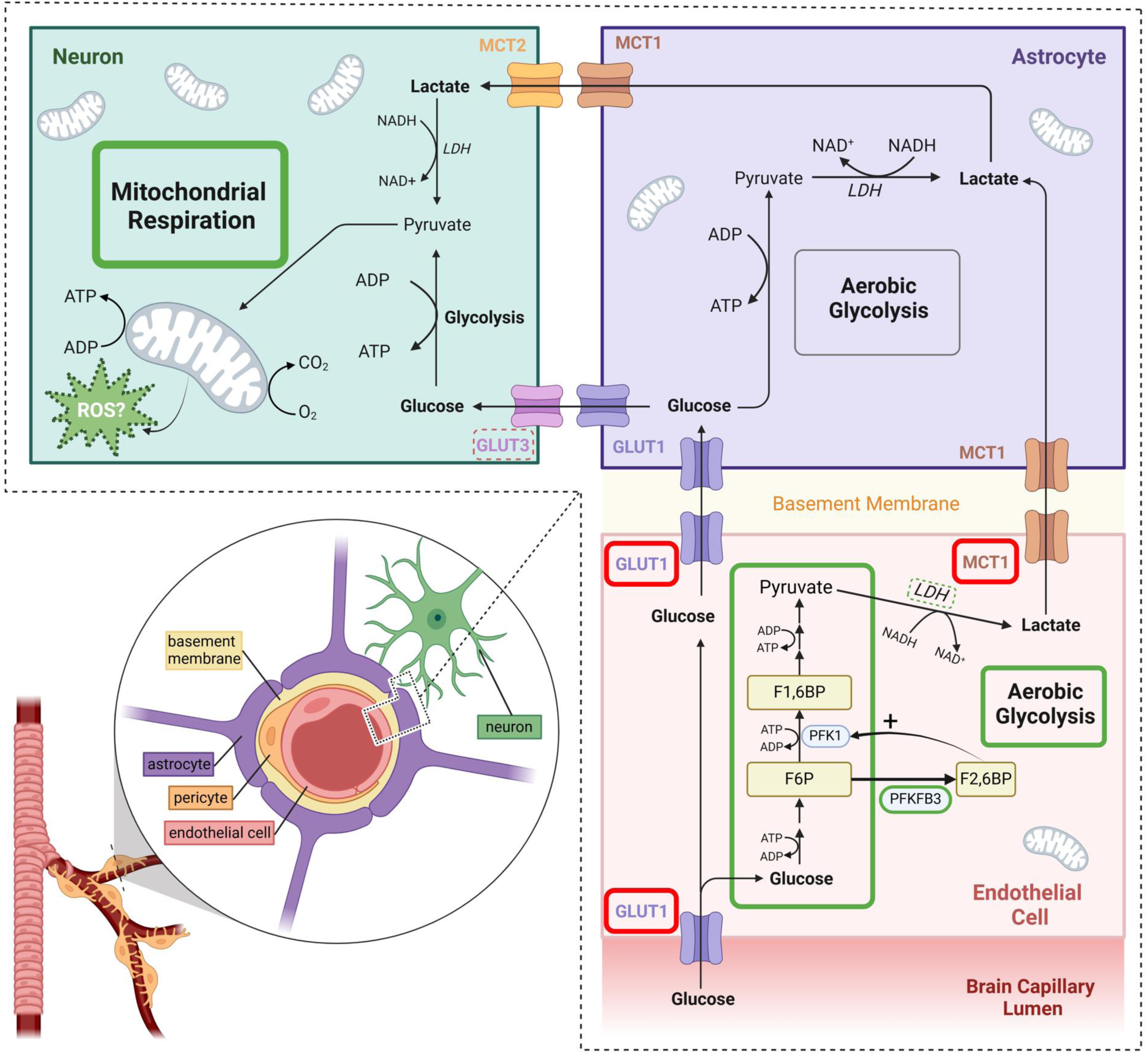
Energy substrate transport and metabolism in the neurovascular unit. <u>Lower left</u>: Depiction of the BBB consisting of specialized endothelial cells (ECs), a basement membrane, pericytes and astrocyte endfeet. Combined with adjacent neurons, the BBB forms the neurovascular unit (NVU). <u>Lower right and top</u>: The transport and metabolism of energy substrates within the NVU is highly dependent on the cell type. Glucose is transported from the brain capillary lumen into ECs by GLUT1 (an ATP- and insulin-independent glucose transporter moving glucose along its gradient across the cell membrane). ECs almost exclusively rely on aerobic glycolysis for ATP production, forming lactate in the process. Any surplus EC glucose is transported towards the basement membrane and into astrocytes (both through GLUT1), where it can either be metabolized through (aerobic) glycolysis into pyruvate and lactate, or be transported into the brain interstitial space, and from there into neurons mainly through neuronal GLUT3. Lactate derived from EC aerobic glycolysis is transported towards the basement membrane through MCT1. Both pericytes (not depicted) and astrocytes transport lactate into the cytoplasm through mainly MCT1. Pericytes are functionally dependent on EC-derived lactate as an energy source^37^. Astrocytes largely rely on glycolysis for ATP production, and can increase their capacity of anaerobic glycolysis on demand, when neuronal activity is high^38^. Excess astrocytic lactate (both derived from ECs and from astrocytes themselves) is transported into the interstitial space through mainly MCT1, where it is transported into neurons by MCT2. This seemingly complex shuttling of energy metabolites, where ECs and astrocyte almost exclusively rely on glycolysis for ATP, ensures a stable neuronal source of both glucose for ‘fast but low-yield’ ATP-production through glycolysis, as well as pyruvate (from transported lactate and from neuronal glycolysis) and oxygen for ‘slow but high-yield’ ATP production in mitochondria. <u>Green and red boxes</u>: In SSD insular cortex BBB, we found a pathway upregulation of glycolysis (**Green box**, lower right) and a (non-significant) upregulation of *LDH*. These findings support the presence of increased aerobic glycolysis in BBB ECs in SSD. Combined with reduced expression of BBB EC *GLUT1* and *MCT1* (**Red boxes**), this suggests a downregulation of *GLUT1* and *MCT1* in reaction to high substrate availability in BBB (i.e., glucose and lactate). Finally, *PFKFB3* expression was significantly increased in SSD BBB. Since *PFKFB3* is the main driver of glycolytic flux in ECs^35^, this increased expression may drive the glycolysis pathway upregulation in SSD BBB. In SSD insular cortex parenchyma, we found an upregulation of mitochondrial energy production pathways (**Green box**, upper left), combined with a (non-significant) downregulation of *GLUT3*. These findings could fit with high energy substrate availability for neurons. A byproduct of mitochondrial respiration is the formation of ROS^39^, a surplus of which contributes to oxidative stress. Increased brain oxidative stress is believed to play a role in the pathophysiology of SSD^40^. However, in this small study, we were unable to detect changes in the oxidative stress response in insular cortex parenchyma. **<u>Abbreviations</u>:** ADP: adenosine diphosphate; ATP: adenose triphosphate; BBB: blood-brain barrier; EC: endothelial cell; F1,6BP: fructose-1,6-bisphosphate; F2,6BP: fructose-2,6-bisphosphate; F6P: fructose-6-phosphate; GLUT1: glucose transporter 1; GLUT3: glucose transporter 3; LDH: lactate dehydrogenase; MCT1: monocarboxylate transporter 1. MCT2: monocarboxylate transporter 2; NVU: neurovascular unit; PFK1: phosphofructokinase-1; PFKFB3: 6-phosphofructo-2-kinase/fructose-2,6 bisphosphatase 3; ROS: reactive oxygen species.

### Decreased Expression of Glucose and Lactate Transporters in SSD Insular BBB

To further elucidate the role of energy substrate transporters in relation to the enriched *Aerobic Glycolysis* pathway in SSD BBB, we specifically examined the gene expression levels of glucose and lactate transporters in the BBB and parenchyma AOIs (**Figure 6**). These transporters include 1) SLC2A1 (GLUT1): the main glucose transporter in BBB ECs, also expressed at low levels in astrocytes^41,42^; 2) SLC2A3 (GLUT3): the main neuronal glucose transporter^43^, also expressed at low levels in BBB ECs^42,44^; 3) SLC16A1 (MCT1), the main lactate transporter in both BBB ECs and astrocytes, as lactate is produced through aerobic glycolysis in these cells^34,35^ (**Figure 5**). Lactate derived from aerobic glycolysis in ECs is transported by MCT1 to both pericytes and astrocytes. Astrocytes, in turn, use MCT1 to transport lactate derived from astrocytic aerobic glycolysis, and from ECs, into the interstitial space^43^. Finally, neurons take up lactate from the interstitial space through SLC16A7 (MCT2) to eventually be used as an energy substrate in mitochondria.

We found a significantly decreased expression of *SLC2A1* (*GLUT1*) and of *SLC16A1* (*MCT1*) in SSD BBB, while no significant change was detected in the expression of *SLC2A3* (*GLUT3*). In contrast, there was no significant difference in the expression of *SLC2A3* (*GLUT3*) or *SLC16A1* (*MCT1*) in parenchyma. Importantly, *SLC16A7* (*MCT2*) did not pass QC and pre-processing. See **Table 3** for statistics and **Figure 6** for a background on energy substrate transport and metabolism in the neurovascular unit.

## Discussion

In this study, we conducted the first spatial profiling analysis investigating the possible role of the insular cortex BBB in the pathophysiology of SSD. To achieve this, we utilized a state-of - the-art spatially resolved transcriptomics technique (GeoMx DSP WTA) to generate whole transcriptome profiles of the BBB and the surrounding brain parenchyma in post-mortem brain tissue of SSD compared to controls. Insular cortex was selected due to its known involvement in SSD^23,24^, and because this region remains largely understudied in post-mortem SSD research. Selecting insular cortex ROIs containing only small-calibre blood vessels, combined with a nuclear marker and an EC marker, and using adjusted settings to include cells bordering on ECs (i.e., pericytes and astrocyte end-feet) technically enabled us to enrich for BBB ECs in the BBB AOI. The enrichment was further improved by excluding areas with aspecific fluorescence in other channels from the BBB AOI (mostly lipofuscin-induced autofluorescence). Using this approach, we were able to include a minimum of 47 cells/AOI for downstream WTA, providing sufficient sensitivity to robustly detect differentially expressed genes with medium and high expression levels, as well as to perform reliable pathways analysis^45^. The study’s robustness was further enhanced by analysing 3 biological replicates per AOI in each sample. In spite of practical and technical challenges (e.g., the globally limited availability of post-mortem brain tissue from SSD patients with a low post-mortem interval, storage duration < 15 years, as well as the relatively low number of cells within the BBB AOI), we found this technique and setup are effective to study the BBB in SSD. Using a commonly accepted log2FC cut-off of ≥ 0.25 for post-mortem psychiatric studies^15^, we identified 265 DEGs in the BBB and 6 in the parenchyma comparing SSD to controls. Cluster analysis showed a clear separation between both BBB and parenchyma, and SSD and control BBB, indicating a distinct BBB transcriptional phenotype in SSD. No such separation was observed in the parenchyma. Results from differential expression and pathway analysis (FGSEA) suggest increased glucose metabolism (glycolysis) in the BBB alongside enhanced mitochondrial energy metabolism in the insular cortex parenchyma of SSD patients. Alterations in brain energy metabolism (both glucose and mitochondrial metabolism) are considered an important pathophysiological model for SSD^6^. To contextualize these findings, we first provide an overview of energy substrate transport and metabolism in the BBB and in brain parenchyma (**Figure 6**).

The adult human brain predominantly relies on glucose as an energy source^43^. Glucose is transported from the brain capillary lumen into BBB ECs mainly by Glucose Transporter-1 (GLUT1; gene: *SLC2A1*). GLUT1 is an insulin-independent glucose transporter that facilitates glucose transport along its gradient across the cell membrane^46^, making intracellular glucose levels in ECs independent of insulin signalling. GLUT1 is highly and selectively expressed in BBB ECs, with much lower expression in glial cells and no expression in neurons^43^.

ECs exhibit a distinct metabolic phenotype: they are highly glycolytic even in the presence of ample oxygen (the ‘Warburg effect’ or ‘aerobic glycolysis’), while normally glycolysis is inhibited under high oxygen conditions due to a shift towards the more ATP-yielding mitochondrial oxidative phosphorylation (‘Pasteur’s effect’)^47^. In aerobic glycolysis, pyruvate is converted into lactate by LDH^46^. Around 85% of EC ATP is generated through aerobic glycolysis. Their glucose consumption rate is remarkably high, rivalling that of cancer cells^34,35^. Contrarily, mitochondria constitute only 2-5% of EC cytoplasmic volume and primarily serve as biosynthetic hubs rather than for ATP production^34^. Furthermore, when glucose levels are elevated, ECs further upregulate aerobic glycolysis while suppressing oxidative phosphorylation, a phenomenon known as the ‘Crabtree effect’^48^.

Similar to ECs, astrocytes primarily rely on aerobic glycolysis rather than oxidative phosphorylation for energy production^49^. This metabolic shift in BBB ECs and astrocytes helps preserve key energy substrates (e.g., lactate) and oxygen for neurons, which depend on the high-ATP-yielding oxidative phosphorylation to sustain their function^34,47,50^. Due to their reliance on mitochondrial respiration, neurons are particularly vulnerable to disruptions in mitochondrial function in all of its facets (‘mitochondrial dysfunction’). For instance, an impaired ability to buffer oxygen radicals generated by oxidative phosphorylation can lead to oxidative stress, contributing to neurodegeneration^51^.

Lactate derived from aerobic glycolysis in BBB ECs and astrocytes is transported through MCT1 (ECs and astrocytes) and MCT4 (astrocytes) into neurons (MCT2) for use in oxidative phosphorylation^52,53^. Residual glucose is transported through GLUT1 (ECs), and GLUT3 (astrocytes) into neurons (GLUT3).

Neurons coordinate their energetic needs with astrocytes and ECs through a process called ‘neurovascular coupling’, the mechanisms of which remain largely unexplored^38^. During heightened neuronal activity, neurons locally increase blood flow to ensure an adequate supply of nutrients and oxygen^3^. BBB ECs play a central role in neurovascular coupling by sensing neural activity through e.g., potassium channels, GABAA- and NMDA-receptors^3^. Furthermore, neurons can induce the Crabtree effect in surrounding astrocytes, thus ensuring transport of sufficient bioenergetic substrates needed to sustain their function^38,54^. As such, metabolism within the BBB appears to be at least partially regulated by the metabolic needs of downstream neurons. Interestingly, this mechanism is reminiscent of tumour ECs, which exhibit a hyperglycolytic phenotype in order to fuel tumour growth^55^.

Glycolytic flux in ECs, astrocytes and neurons is mainly regulated through PFKFB3, an enzyme converting F6P into F2,6BP. In turn, F2,6BP is a potent allosteric activator of phosphofructokinase-1 (PFK1), the rate-limiting enzyme of glycolysis (**Figure 5** and **6**)^34^. Both ECs and astrocytes express high levels of PFKFB3 to sustain glycolytic flux^56^. PFKFB3 acts as the principal inducer of glycolysis in both astrocytes^56^ and ECs^35,55^, particularly during high bioenergetic demand, such as in hypoxia, physiological angiogenesis and tumour angiogenesis. In contrast, neurons maintain a low glycolytic flux by actively degrading PFKFB3^50,57^.

This background on energy substrate transport and metabolism in the NVU provides context for our findings. We observed an enrichment in *Glycolysis* pathway genes, a significant increase in *PFKFB3* expression and a (non-significant) increase in *LDH* expression in SSD BBB compared to controls. These findings suggest an upregulation of aerobic glycolysis in SSD BBB via *PFKFB3* induction. Additionally, we found a significantly decrease in *GLUT1* and *MCT1* expression in SSD BBB, without a compensatory upregulation of *GLUT3*. Given the concurrent increase in aerobic glycolysis gene expression, this downregulation may reflect a response to high substrate availability in the BBB (glucose, lactate). Some studies suggest that *GLUT1* expression in EC inversely correlates with blood glucose levels (i.e., downregulated in hyperglycaemia and upregulated in hypoglycaemia), though this remains debated^46,58^. Ample bioenergetic substrate availability is also supported by our finding of increased expression of mitochondrial energy metabolism pathway genes and a significant increase in *COX7A1* expression in SSD insular cortex parenchyma. An impairment of the EC/astrocyte-to-neuron glucose and lactate shuttle as an alternative explanation to reduced levels of BBB *GLUT1* and *MCT1* would lead to decreased parenchymal mitochondrial respiration in the parenchyma, which we did not observe. To our knowledge, decreased *GLUT1* expression in SSD BBB has not been previously described. Sullivan et al. found reduced *GLUT1* and *GLUT3* expression, increased *MCT1* and decreased glycolytic enzyme expression (*HK* and *PFK1*) in laser-capture microdissected (LCM) pyramidal neurons (but not astrocytes) in the dorsolateral prefrontal cortex (DLPC) in SSD. However, they did not investigate the BBB or the insular cortex^59^.

While speculative at this stage, our findings could point towards increased mitochondrial energy metabolism in the insular cortex parenchyma in SSD, supplied by a heightened glycolytic flux in the BBB induced by upregulated expression of *PFKFB3*. Alternatively, elevated BBB glycolysis may actively drive mitochondrial energy metabolism in the insular cortex in SSD. Notably, fMRI studies have previously reported insular cortex hyperactivity in SSD patients in response to neutral stimuli^27^.

Recent hypotheses propose that cerebral hypermetabolism induced by (transient) hyperglycolysis could contribute to bipolar mania^60^. Since SSD and bipolar disorders have overlapping etiologies^61^, a hypermetabolic state in both the BBB and insular cortex parenchyma may represent a disease-driving mechanism in SSD. If so, targeting hyperglycolysis in the BBB could offer a novel therapeutic strategy to restore brain parenchymal metabolic balance in SSD. Additional explanations for our findings could include mitochondrial dysfunction in SSD insular parenchyma, potentially leading to oxidative stress and neuronal dysfunction (in line with the ‘oxidative stress’ hypothesis of SSD)^62^.

Whether the observed increase in aerobic glycolysis and decrease in *GLUT1* and *MCT1* expression in the SSD insular cortex BBB is part of the primary disease process or a consequence of long-term antipsychotic (AP) treatment remains speculative. Frequently used APs are known to elevate fasting blood glucose level^63^, though the extent varies between different APs^64^. There is some evidence that APs can influence the expression of glycolytic enzymes, including *PFKFB3*, but findings are inconsistent. This variability is likely due to differences in model systems used (both *in vivo* and *in vitro)*, with many studies focussing on neurons, which as explained have inherently different energy metabolism properties than BBB ECs and astrocytes^65–68^. Since *PFKFB3* expression in ECs can be induced by various different stimuli, such as circulating growth factors, hormones, hypoxia^34–36,69,70^, the current study is not designed to determine the precise mechanism underlying its increased expression.

Alterations in glycolysis and mitochondrial energy metabolism have been previously reported in post-mortem SSD brain transcriptomic and proteomic studies^6^. However, the majority of these studies have found a reduced expression of genes and proteins involved in glycolysis and mitochondrial energy metabolism^6,71–73^. Combined with an intrinsic association of SSD with dysregulated glucose metabolism, such as a genetic predisposition to insulin resistance, hyperglycaemia and type 2 diabetes, these findings gave rise to the hypothesis that a cerebral bioenergetic deficiency may play a causative role in the pathophysiology of SSD^6,74^. Our findings of increased BBB aerobic glycolysis and increased parenchyma mitochondrial metabolism gene expression in SSD insular cortex are in apparent contrast with these prior findings. A key reason for this discrepancy may be the predominant focus of prior research on the DLPC, while other brain regions remain largely understudied. Although both the DLPC and insular cortex are implicated in SSD, their functions are incomparable, playing roles in e.g., executive functioning versus interoception/(self) awareness, respectively^23,24,75^. Furthermore, studies indicate considerable regional variation in BBB function^3^, meaning molecular findings in one brain region cannot be extrapolated to another. Indeed, our results align with the only other post-mortem study on the insular cortex in SSD: Pennington et al. used LCM to analyse insular cortex layer 2 (i.e., containing a mixture of neurons, glial cells, BBB cells) and found increased protein levels of glycolytic enzymes (ALDOC) and of a mitochondrial ATPase subunit (ATP5C1)^76^. Notably, their study used tissue from the Stanley Brain Foundation, the same tissue repository used in other studies reporting decreased glycolysis and mitochondrial energy metabolism in DLPFC. These contrasting findings in different brain regions derived from material from the same donor cohort suggest that SSD may involve region-specific metabolic alterations rather than a generalized bioenergetic deficiency as previously proposed^6^.

Another possible explanation is that post-mortem transcriptomic and proteomic studies in SSD rely on tissue from a limited number of brain banks, primarily the Stanley Brain Collection, which may restrict the generalizability of findings. Furthermore, these studies often involve material with a long PMD and low tissue pH, both of which can significantly impact mitochondrial gene expression^29,77^. In our sample set, only one SSD sample exhibited significant agonal changes and a cerebrospinal fluid (CSF) pH <6 (5.93). All other samples had a pH>6 and did not display (post-)agonal changes on neuropathological examination. To ensure this sample did not unduly influence our finding, we repeated the entire data analysis excluding it. The results remained unchanged, confirming the robustness of our conclusion (data not shown).

A crucial distinction of our study is the use of a spatially resolved approach, allowing us to analyse the BBB and brain parenchyma separately. Most prior studies relied on homogenized brain tissue, containing a mixture of e.g., neurons, glial cells and BBB cells. Because of the distinct metabolic phenotype of these cell types (**Figure 6**), analysing bulk tissue may obscure critical metabolic differences. Some studies attempted to enrich specific cells using LCM, primarily focusing on DLPC neurons^59^. One small study applied LCM to isolate microvascular BBB from DLPC of SSD donors and controls for transcriptomic analysis, reporting a hypo-inflammatory BBB phenotype in SSD but no metabolic alterations^78^. Differences in study design, including the number of studied samples, the use of LCM to enrich for BBB, cDNA conversion and microarray for transcriptomic analysis, frozen DLPC tissue, and a longer PMD may explain the lack of metabolic findings in that study compared to ours. Another study employed single-nucleus RNA sequencing on sorted cell types, including ECs, from dissociated SSD midbrain tissue compared to controls^7^, and did not find DEGs in ECs. However, the absence of DEGs may stem from the use of a non-spatially resolved technique, the limited number of microvascular ECs sequenced, and the selection of midbrain, an anatomically complex region with diverse nuclei and functions. Notably, that study did identify DEGs in ependymal cells (which form the blood-cerebrospinal fluid barrier, not the BBB) and pericytes. Taken together, these methodological differences likely explain the discrepancies between previous studies and our findings.

Our study is the first to apply spatial WTA to SSD insular cortex tissue, enabling the separate analysis of the BBB and parenchyma. While the inclusion of 16 samples for spatial biology studies is substantial, our findings cannot be generalized to the entire SSD spectrum. In that context, we consider the relatively small sample size as one of the potential limitations of this study. In addition, solely profiling the insular cortex restricts the detection of potentially contrasting metabolic signatures in other brain regions.

In conclusion, using spatial transcriptomics (GeoMx DSP WTA), we identified a hyperglycolytic transcriptional phenotype in the BBB and increased mitochondrial energy metabolism in the surrounding parenchyma in insular cortex of SSD patients. These findings add nuance to the existing understanding of brain bioenergetic alterations in SSD, suggesting that metabolic changes may be region-specific rather than generalized. Our results highlight the need for a broader ‘brain mapping’ approach examining multiple brain regions from the same donor. Additionally, the distinct metabolic profiles of the BBB and brain parenchyma emphasize the importance of spatial multi-omics in post-mortem psychiatric research and the potential for therapies targeting BBB function in SSD.

